# Comparative transcriptomics identifies *Botrytis cinerea* induced conserved defences across different tissues of *Fragaria vesca*

**DOI:** 10.1101/2020.07.18.210138

**Authors:** Raghuram Badmi

## Abstract

Grey mould is one of the most devastating plant diseases that causes huge losses to the agricultural sector worldwide. *Botrytis cinerea*, the causal pathogen of grey mould, is a generalist necrotrophic fungus with the ability to infect over 1000 species and influence a broad array of host’s physiological responses. *B. cinerea* is capable of infecting a wide variety of tissues such as leaves, stem, flowers and fruits that adds to the already complex problem in controlling its spread. Understanding the pathogen induced transcriptional reprogramming in different tissues is important to identify key genes for targeted gene modifications. Identifying the genes that are common between different tissue infections will reveal similarities and differences between these pathogen-tissue pairs. In this study, the transcriptomic datasets of Botrytis infected white berries of *Fragaria vesca* (*WhiteBc*) and Botrytis infected red berries of *F. vesca* (*RedBc*) were (re)mapped to the latest *F. vesca* transcriptome to enable direct comparisons with the Botrytis infected *F. vesca* leaves (*LeafBc*). The genes involved in MAP kinase signalling, pathogenesis-related, allergens, cell-wall defences, detoxification and secondary metabolites were <u>Co</u>mmon <u>Re</u>sponsive and <u>Up</u>regulated (CoReUp) between *LeafBc*, *WhiteBc* and *RedBc*, suggesting their important roles against *B. cinerea* infection in all three tissues. These insights maybe helpful for generating *B. cinerea* resistant varieties of strawberry.

## Introduction

The necrotrophic fungi *Botrytis cinerea* is one of the most devastating pathogens that is capable of causing a loss of over €1 billion per year (Dean et al., 2012; Veloso & van Kan, 2018). It causes grey mould disease in more than 1000 plant species that include economically important crops. Some of the high-value crops that suffer severe losses by *B. cinerea* include strawberry, tomato, grapes, raspberry, blackberry, kiwifruit, apples and pears. The wide host range of the fungi is accompanied by wide range of symptoms that are not always similar between organs and tissues. Some of the symptoms from *B. cinerea* infection include soft rotten tissue such as on strawberries and leaves, minute pock marks and blemished lesions on flower petals and rotting on stems (Williamson et al., 2007). Due to its broad host range and infectious nature, it is recognised as a generalist plant pathogen rather than a specialist. Upon infection, *B. cinerea* manipulates the host’s physiological processes for its own benefit to spread infection (Veloso & van Kan, 2018). This is achieved by deploying a wide variety of virulence factors into the host including smallRNAs, effector proteins, toxins, cell-wall degrading enzymes, oxalic acid and proteins that probably induces host programmed cell death (PCD) (Petrasch et al., 2019; Veloso & van Kan, 2018). Infection by *B. cinerea* involves pathogen invasion into the tissues followed by necrosis formation that gradually spreads across the tissues thereby destroying the entire plant. Its necrotrophic lifestyle, broad-host range and generalist nature adds to its devastating effects on infected crops.

Strawberry (*Fragaria × ananassa*), is a high-value crop that suffers severe losses majorly due to postharvest infection worldwide. It is recognised that strawberry is susceptible to infection from more than 50 different fungal genera, making it the most important risk factor in strawberry production (Garrido et al., 2011). Strawberry also suffers both pre- and post-harvest losses due to grey mould disease caused by *B. cinerea*, which infects different tissues of strawberry such as leaves, flowers and berries (Petrasch et al., 2019). Although, berries are the commercially important parts of strawberry plant, infection on leaves might weaken the plant by shifting plant’s resources to defence, thereby yielding low quality fruits. Infection on flowers would already reduce the crop yield by destroying the berry development itself. Understanding the transcriptomic responses to *B. cinerea* infection in different tissues of strawberry will reveal important insights that could be crucial to understand this plant-pathogen interaction pair. For crop improvement programs, it can be promising to rely on genes that are commonly involved in defence responses in various tissues as compared to the genes responsive in only one tissue, given that the pathogen is capable of infecting different tissues simultaneously.

Several studies reported transcriptomic responses of *Fragaria vesca*, a diploid model for commercial strawberry (*F. ananassa*), upon *B. cinerea* infection. *B. cinerea* infected leaves of *F. vesca* display large transcriptomic changes that include about 7000 differentially expressed genes (Badmi et al., 2019). Transcriptomic responses of *F. vesca* berries were studied upon *B. cinerea* infection to understand the low susceptibility of unripe (white) fruits as compared to ripe (red) fruits. Comparison between the *B. cinerea* responsive genes in unripe and ripe berries revealed important differences that pointed out to the susceptible nature of ripe red berries due to their softened cell-walls (Haile et al., 2019). The authors reasoned that upregulation of cell-wall strengthening genes and downregulation of cell-wall softening genes might indicate the relatively higher resistance of white unripe berries to *B. cinerea* (Haile et al., 2019). This points out that the quantitative resistance against *B. cinerea* may be tissue dependent and reflects its specific physiology. Building upon this observation, comparison of transcriptomic responses between different tissues will discover commonly regulated genes across tissues against *B. cinerea* infection. Such a comparison might shed light on the important genes that are active across tissues that are yet unidentified to confer resistance against *B. cinerea*. Furthermore, the insights might reveal novel gene targets that will be crucial in crop improvement breeding programs.

## Materials and Methods

### RNA-seq data retrieval and processing

The raw RNA-seq read files (SRA files) were downloaded in windows command line terminal using ‘fastq-dump’ command. Fastp (Chen et al., 2018) was used for quality control, adapter trimming and quality filtering. For the alignment of resulting reads to the latest *F. vesca* transcriptome (Edger et al., 2018) (ftp://ftp.bioinfo.wsu.edu/species/Fragaria_vesca/Fvesca-genome.v4.0.a1) rsem aligner with bowtie2 (Li & Dewey, 2011) option was used. From the gene results files, fragments per kilobase of exon per million reads mapped (FPKM) values were extracted and EdgeR (McCarthy et al., 2012; Robinson et al., 2009) was used for differential expression analysis. For normalization, the ‘trimmed mean of M values’ (TMM) method was used and the p-values were adjusted using Benjamini and Hochberg approach (Benjamini & Hochberg, 1995).

### Analysing differentially expressed genes

The differentially expressed genes from three tissues of *F. vesca* – Botrytis interactions (*LeafBc*, *WhiteBc* and *RedBc*) were arranged to have the same *F. vesca*_4.0 IDs and retained the ones with log2 fold change (log2FC) expression values > 1.5 and p-adjusted values < 0.05. The lists of up- and down-regulated genes were separated and used to generate venn diagrams using InteractiVenn online tool (Heberle et al., 2015). The list of genes intersecting between the pairs were extracted by clicking on the intersecting area of the venn diagram. Matplotlib tool (https://matplotlib.org/) in Python 3.6 was used to visualize the 3D scatter plot of log2FC values between the three sets.

## Results and Discussion

### *Botrytis cinerea* induced transcriptional reprogramming in three *Fragaria vesca* tissues

Mapping the RNA-seq datasets to a single reference transcriptome will provide common gene IDs between studies whose expression values can be directly compared between each other. The transcriptome data from *B. cinerea* infected leaves (*LeafBc*) is mapped to the *F. vesca*_4.0.a1 transcriptome (Badmi et al., 2019). To enable reliable comparisons across studies, the transcriptomic datasets of white berry infected with *B. cinerea* (*WhiteBc*) and red berry infected with *B. cinerea* (*RedBc*) (Haile et al., 2019) were mapped to the same *F. vesca*_4.0.a1 transcriptome (Edger et al., 2018). Furthermore, as the transcriptome datasets of these three tissues are at 24-hour post infection, comparative studies between them will provide streamlined insights. Table 1 lists the SRA accession numbers of the transcriptome datasets used for mapping and analysis. Table S1 lists all the 28588 *F. vesca* gene IDs with the corresponding log2FC and p-values in the three *F. vesca* tissues against *B. cinerea* infection. The log2FC values with a p-value of less than 0.05 (p < 0.05) were retained after filtering and used for comparisons between three different tissue infections. In *LeafBc* tissues, 3689 genes were upregulated and 3314 genes were downregulated constituting about 11% and 9% of the transcriptome respectively (Badmi et al., 2019). In *WhiteBc*, 526 genes were upregulated and 476 genes were downregulated constituting about 2% and 1% of the transcriptome respectively. In *RedBc*, 408 genes were upregulated and 634 genes were downregulated constituting about 1% and 2% of the transcriptome respectively (Fig. 1, Table S2). For *WhiteBc* and *RedBc*, the number of up- and downregulated genes were different in this study as compared to the previous study (Haile et al., 2019), probably because of the difference in the transcriptome reference used. As a validation step before in-depth analysis, the expression values from 15 randomly selected genes were compared between this study and from Haile et al., (2019). As shown in Table 2, the expression values between studies are proportional and comparable with each other, thereby ascertaining the reliability of the methods used in this study.

**Table 1:** Details of the genotype, tissue, samples and their SRA accession numbers used for mapping the RNA-seq reads to the *F. vesca* transcriptome and their corresponding number of differentially expressed genes.

| Pathogen | Tissue | RNA-seq sample | NCBI SRA accession numbers | Upregulated<br>log2FC > 1.5 | Downregulated<br>log2FC > 1.5 | P<br>adjusted | Reference |
| --- | --- | --- | --- | --- | --- | --- | --- |
| *Botrytis cinerea* | Leaf | n.a. | n.a. | 1535 | 1373 | < 0.05 | (Badmi et al., 2019) |
| *Botrytis cinerea* | White Berry | ControlWhiteFruit24hpi_rep1 | SRR8840923 | 526 | 476 | < 0.05 | (Haile et al., 2019) |
|  |  | ControlWhiteFruit24hpi_rep2 | SRR8840922 |  |  |  |  |
|  |  | ControlWhiteFruit24hpi_rep3 | SRR8840917 |  |  |  |  |
|  |  | Botrytis inoculated WhiteFruit24 hpi_rep1 | SRR8840916 |  |  |  |  |
|  |  | Botrytis inoculated WhiteFruit24 hpi_rep2 | SRR8840919 |  |  |  |  |
|  |  | Botrytis inoculated WhiteFruit24 hpi_rep3 | SRR8840918 |  |  |  |  |
| *Botrytis cinerea* | Red Berry | ControlRedFruit24hpi_rep1 | SRR8840921 | 408 | 634 | < 0.05 | (Haile et al., 2019) |
|  |  | ControlRedFruit24hpi_rep2 | SRR8840920 |  |  |  |  |
|  |  | ControlRedFruit24hpi_rep3 | SRR8840927 |  |  |  |  |
|  |  | Botrytis inoculated RedFruit24 hpi_rep1 | SRR8840926 |  |  |  |  |
|  |  | Botrytis inoculated RedFruit24 hpi_rep2 | SRR8840925 |  |  |  |  |
|  |  | Botrytis inoculated RedFruit24 hpi_rep3 | SRR8840924 |  |  |  |  |

**Table 2:** List of gene expression values from Haile et al., (2019) and this study, for comparison of the expression values between the studies. This table serves as a validation for the reliability of expression values of the genes across studies and across two different transcriptome references.

| *F. vesca*_4.0_ID | Phytozome ID | log2FC from Haile et al., 2019 |  | log2FC from this study |  |
| --- | --- | --- | --- | --- | --- |
|  |  | White berry | Red berry | White berry | Red berry |
| FvH4_1g28000.1 | gene21072 | 1.210 | n.a. | 1.033 | 0.287 |
| FvH4_6g22790.1 | gene00522 | 1.797 | 1.746 | 2.045 | 1.762 |
| FvH4_4g18850.1 | gene07064 | 2.524 | 1.531 | 2.651 | 1.383 |
| FvH4_3g42760.1 | gene20547 | 1.201 | 2.123 | 0.605 | 1.267 |
| FvH4_4g36900.1 | gene00814 | -1.055 | -1.003 | -0.862 | -0.861 |
| FvH4_7g16390.1 | gene19196 | -1.112 | 1.506 | -0.358 | 0.587 |
| FvH4_7g29190.1 | gene13301 | 0.908 | -1.035 | 0.768 | -1.160 |
| FvH4_3g19870.1 | gene17154 | -4.427 | 1.874 | -1.682 | 0.990 |
| FvH4_1g15530.1 | gene24085 | 1.757 | 1.263 | 1.991 | 1.029 |
| FvH4_6g03250.1 | gene16534 | -1.009 | 1.132 | -0.850 | 1.154 |
| FvH4_6g47350.1 | gene04330 | -1.815 | 2.749 | -1.340 | 1.626 |
| FvH4_4g13500.1 | gene06526 | -2.772 | 2.594 | -1.103 | 0.654 |
| FvH4_7g28970.1 | gene13326 | -2.174 | 2.348 | -1.602 | 1.398 |
| FvH4_3g08220.1 | gene29796 | 2.361 | 2.209 | 0.452 | 1.456 |
| FvH4_3g28040.1 | gene14626 | 1.714 | 2.167 | 1.570 | 1.994 |

**Figure 1:**
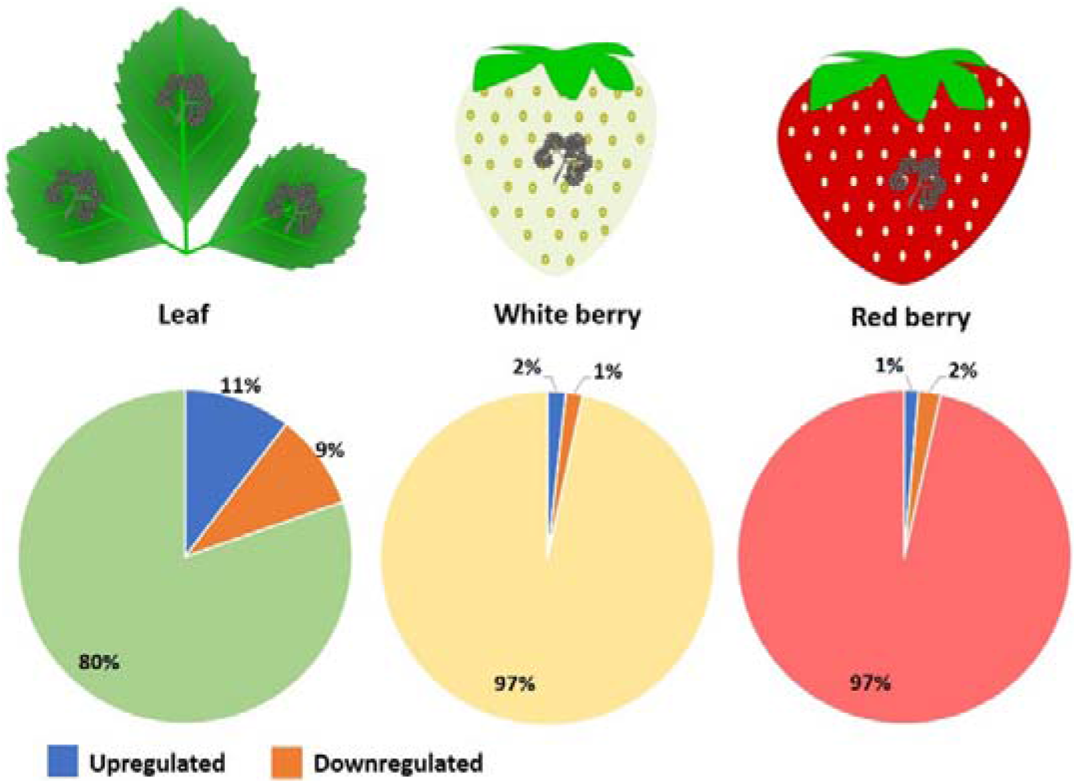
Percentage of *Fragaria vesca* transcriptome responsive to *Botrytis cinerea* infection. Schematic representation of Leaf, White berry and Red berry tissues infected with *B. cinerea* with corresponding venn diagrams depicting the amount of up- and downregulated genes as percentage of the total *Fragaria vesca* transcriptome.

### <u>Co</u>mmon <u>Re</u>sponsive (CoRe) genes of *Fragaria vesca* upon *Botrytis cinerea* infection

*Botrytis cinerea* naturally infects important tissues in strawberry plant including leaves, flowers and berries. Although berries are the economically important and valuable, *B. cinerea* infection on flowers will already lead to decrease in yield, further risking the infection spread to the entire plant. Also, infection on leaves will not only serve as a hotspot for disease spread but also might weaken the plant thereby reducing the resources for a complete development of fruit. Therefore, it is of utmost importance to understand the induced defences and the responsive genes in these tissues that serve as natural hosts for *B. cinerea* infection. Also, comparing the defence responses between these tissues will identify common genes that are responsive to *B. cinerea* infection in these three tissues. To get an overview of the overlapping genes, venn diagram was generated using the filtered (p < 0.05) lists of gene expression values from three Botrytis infected tissues corresponding to 7003 *LeafBc*, 1002 *WhiteBc* and 1042 *RedBc* genes (Table S2). As shown in Fig. 2a, 84 genes are common between these three tissues and are referred to as the <u>Co</u>mmon <u>Re</u>sponsive (CoRe) set of genes against *B. cinerea*.

**Figure 2:**
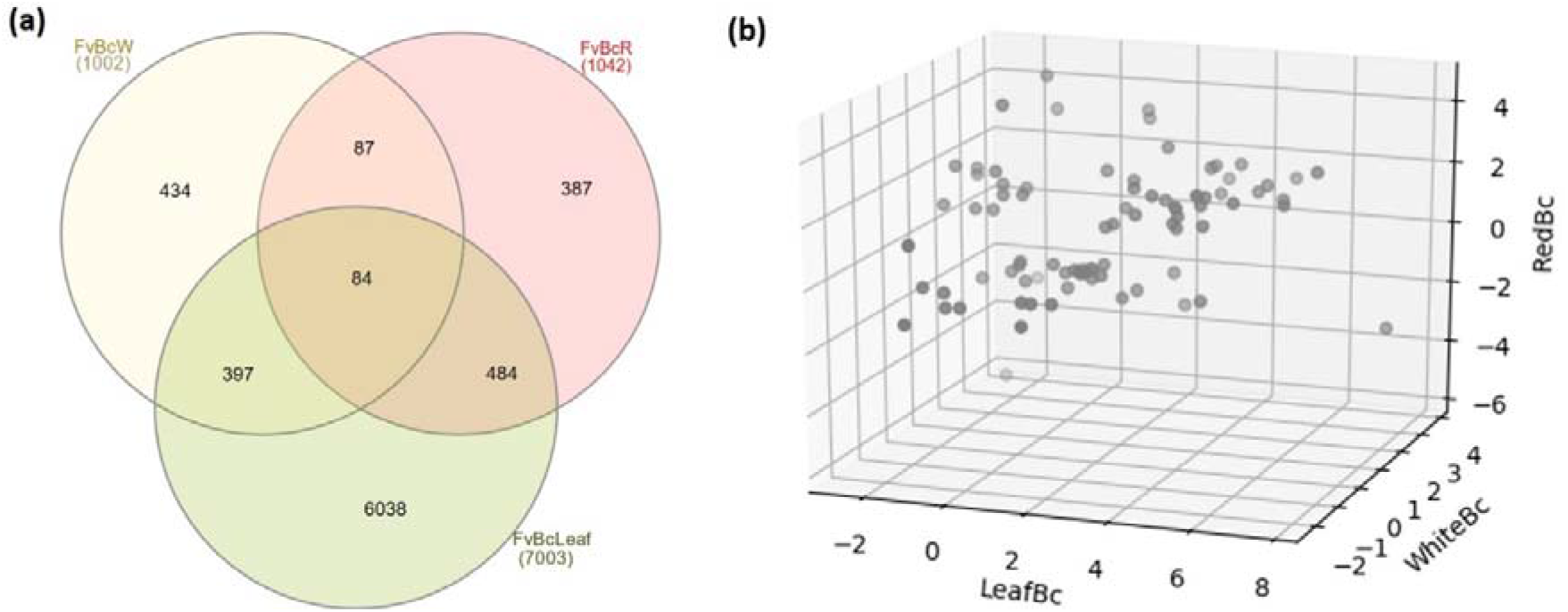
(a) Venn diagram showing numbers of overlapping and specific genes between *F. vesca* – *B. cinerea* interactions in three different tissues and (b) A 3D scatter plot representing 84 Common Responsive (CoRe) set of genes in three tissues of *F. vesca* against *B. cinerea* infection. The three *F. vesca* – *B. cinerea* pairs *LeafBc*, *RedBc* and *WhiteBc* form X-, Y- and Z- axis respectively. Each dot representing each gene is positioned relative to its expression levels in the three tissues of *F. vesca* – *B. cinerea* pairs.

These genes include both up- and downregulated genes in three tissues of *F. vesca* – *B. cinerea* interactions. To visualize the gene expression patterns in three Botrytis infected tissues a 3D scatter plot was generated from the 84 CoRe genes using MatPlotLib (Fig. 2b). Each dot in the 3D scatter plot represents a single gene that is positioned according to its expression levels in each of the three Botrytis infected tissues. Further, separate lists of up- and downregulated genes were used to generate venn diagram and to identify overlapping genes between tissue infections. Several genes were up- or downregulated in a tissue specific manner. Specific upregulation of 3149 genes in *LeafBc*, 205 genes in *WhiteBc* and 146 genes in *RedBc* was observed. Also, specific downregulation of 2889 genes in *LeafBc*, 229 genes in *WhiteBc* and 241 genes in *RedBc* was observed (Fig. 3). Between *LeafBc* and *WhiteBc* 106 genes were commonly upregulated and 167 genes were commonly downregulated. Between *LeafBc* and *RedBc* 119 genes were commonly upregulated and 96 genes were commonly downregulated. A total of 34 genes were <u>Co</u>mmon <u>Re</u>sponsive <u>Up</u>regulated (CoReUp) and 5 were <u>Co</u>mmon <u>Re</u>sponsive <u>Do</u>wnregulated (CoReDown) genes across the three Botrytis infected tissues (Fig. 3), indicating the common defences between these tissues against Botrytis infection. The CoReUp genes offer interesting insights into the conserved modules of defence against *B. cinerea* in different tissues. The 34 CoReUp genes between three Botrytis infected tissues include well-known defence responsive genes such as MAPK signalling component, pathogenesis-related proteins, cell-wall defence proteins, detoxification proteins and secondary metabolites (Table S3, Fig. 4). All of these genes are consistently found to be important in *F. vesca* defences (Amil-Ruiz et al., 2011, 2016; Badmi & Sheikh, 2020), reinforcing their importance against *B. cinerea* infection in strawberry. The polygalacturonase-inhibiting protein (PGIP) that blocks fungal polygalacturonase enzymes from degrading polygalacturonan components of plant cell-walls (Mehli et al., 2004) is one of the CoReUp genes, suggesting its important resistance functions against *B. cinerea*. Chitinase (Vellicce et al., 2006) and β-1,3-glucanase (BG) enzymes that act on fungal cell-walls (Amil-Ruiz et al., 2011) to release highly active elicitor molecules are also upregulated in the CoRe set. Signalling components such as mitogen activated protein kinase (MAP kinase) genes and pathogenesis related proteins such as PR4 and allergens are also upregulated in the CoRe set, thus reinforcing their importance against *B. cinerea* infection in all three strawberry tissues. Furthermore, several genes involved in the biosynthesis of flavonoids, phenolics and phytoalexins are also upregulated in the CoRe set. Induced expression of chalcone isomerase (CHI) and chalcone synthase (CHS) correlates with increased resistance against pathogens and pests probably because of their known functions in biosynthesis of flavonoids (Dowd et al., 2018; Fofana et al., 2002; Pettinga et al., 2018; Zhou et al., 2018). Flavanone 3-hydroxylase which also functions in flavonoid biosynthetic pathway has coordinated expression with CHI and CHS (Pelletier & Shirley, 1996), also pointing out its functions in disease resistance. Phenylalanine ammonia lyase which is an important enzyme in the biosynthesis of salicylic acid and flavonoids (Shine et al., 2016) is also a CoReUp gene. It is interesting to note that multiple genes involved in the detoxification functions such as glutathione S-transferase-like (GST-like) and detoxification proteins were upregulated in the CoRe set (Table S3). GST conjugates xenobiotic compounds such as atrazine and 1-chloro-2,4-dinitrobenzene (CDNB) to the tripeptide glutathione (GSH, γ-L-glutamyl-L-cysteinyl-glycine) thereby detoxifying these toxic compounds (Gullner et al., 2018; Schröder et al., 2007). However, calmodulin-like protein that is involved in cell-wall biosynthesis (Badmi et al., 2018) is upregulated in *LeafBc* and *WhiteBc* and downregulated in *RedBc* (Table S3). Similar response is also seen in case of zinc finger AN1 protein, 60S ribosomal protein and heat shock proteins (Table S3). These observations offer new insights into the common defence mechanisms of strawberry tissues against *B. cinerea* that maybe valuable for generating resistant crop varieties.

**Figure 3:**
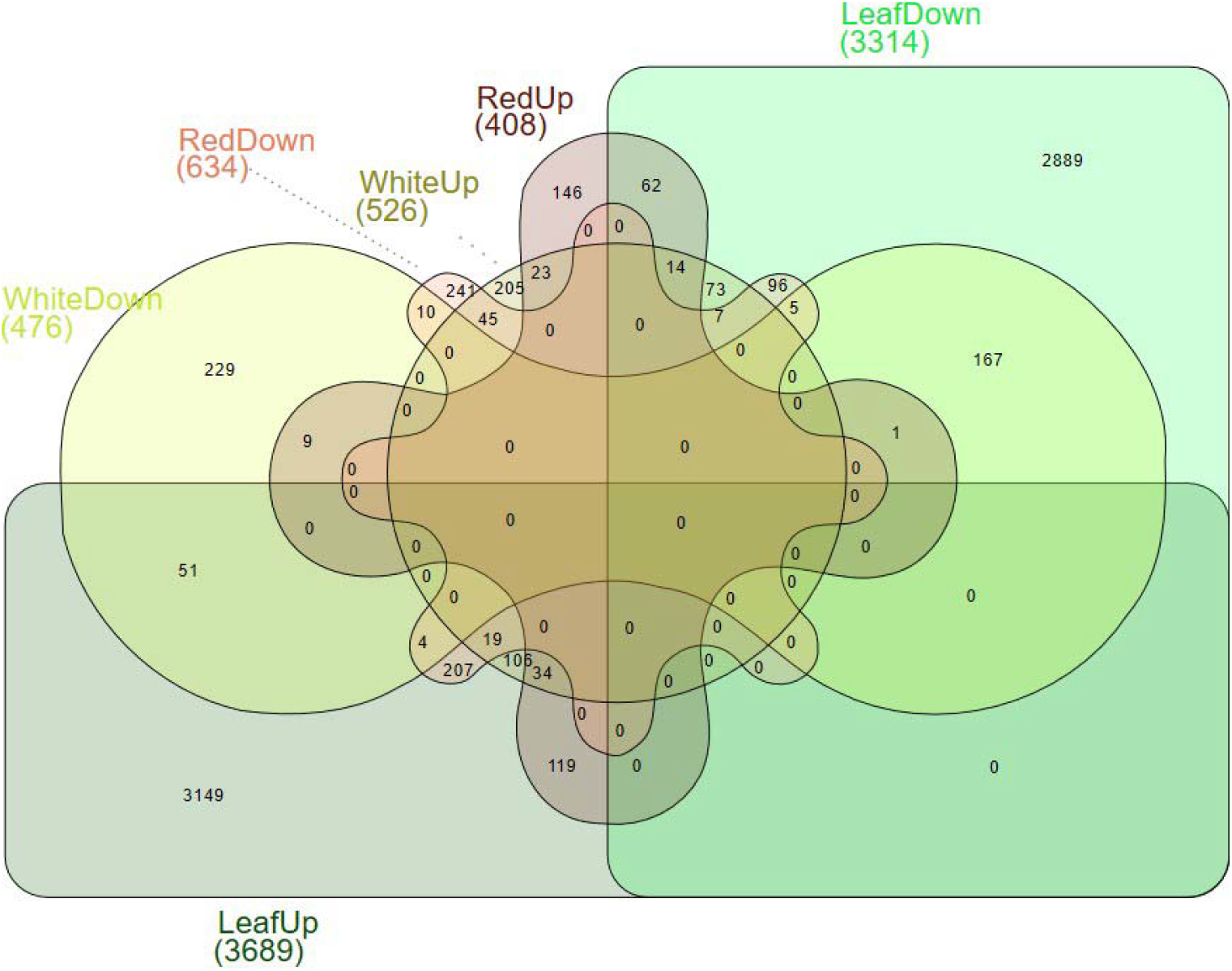
Venn diagram of upregulated and downregulated lists of genes in *F. vesca* – *B. cinerea* interactions in three different tissues showing the overlapping and specific genes in each tissue. The numbers in parenthesis indicate the total number of genes in respective groups.

**Figure 4:**
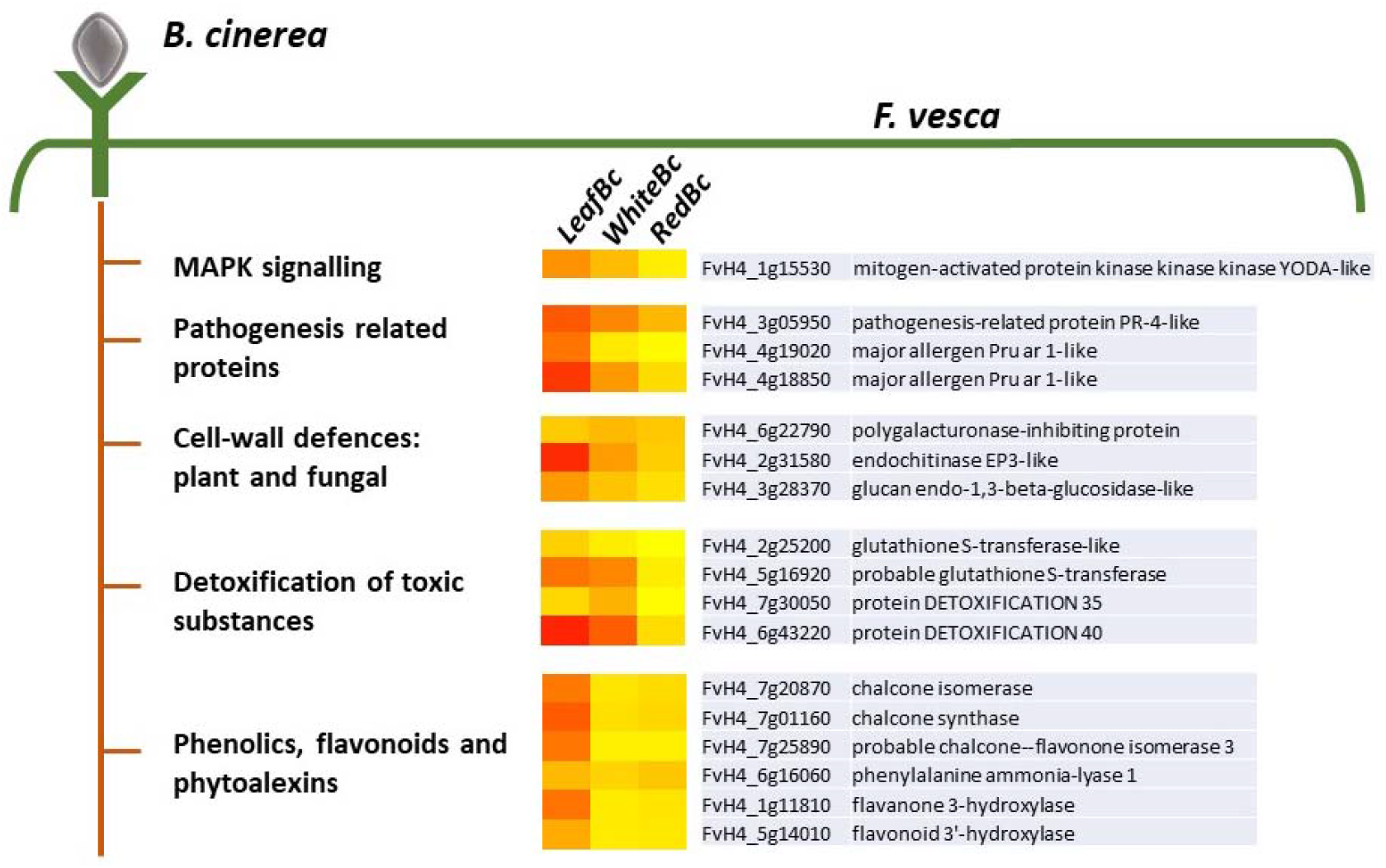
Schematic heatmap representation of the Common Responsive Upregulated (CoReUp) genes in the three *F. vesca* – *B. cinerea* tissues. The expression values of genes are represented as heat maps corresponding tissues and grouped according to their annotated functions.

### Defence responsive genes across different tissues and pathogens in *Fragaria vesca*

The CoReUp genes in *F. vesca* – *B. cinerea* interactions probably reflect their central role against *B. cinerea* infection independent of the tissue types. Furthermore, the availability of the upregulated list of genes against three different pathogen infections in *F. vesca* (Badmi & Sheikh, 2020) allows for comparison with CoReUp genes in this study. A total of 13 genes were found to be common between CoReUp list in this study and ‘Upregulated in all *FvPathogen*’ list from Badmi and Sheikh (2020) (Table 3). It is worth noting that ‘Upregulated in all *FvPathogen*’ list (Badmi & Sheikh, 2020) includes three most important strawberry pathogens with different lifestyles – necrotrophic *B. cinerea* infected leaves, hemibiotrophic *Phytophthora cactorum* infected roots and biotrophic *Podosphaera aphanis* infected leaves. Therefore, the 13 upregulated genes (Table 3) encompasses three different pathogen infections and different tissues such as leaves, roots, white berry and red berry, pointing out that these genes can be central for broad-spectrum disease resistance in different tissues of *F. vesca*. The most prominent genes such as endochitinase, PR4, BG, allergens, GST-like and PGIP are found in this list thus reinforcing their importance in broad spectrum disease resistance and in different tissues of *F. vesca*. However, there were no overlapping genes between the downregulated lists from these two studies. These insights reflect that the 13 commonly upregulated genes across tissue infections and different pathogens might have similar pathogen responsive cis-elements in their promoters that drive their upregulation upon infection. Furthermore, follow-up studies are required to investigate if the gene upregulations translate into protein functions and/or corresponding defence metabolite accumulations.

**Table 3:**
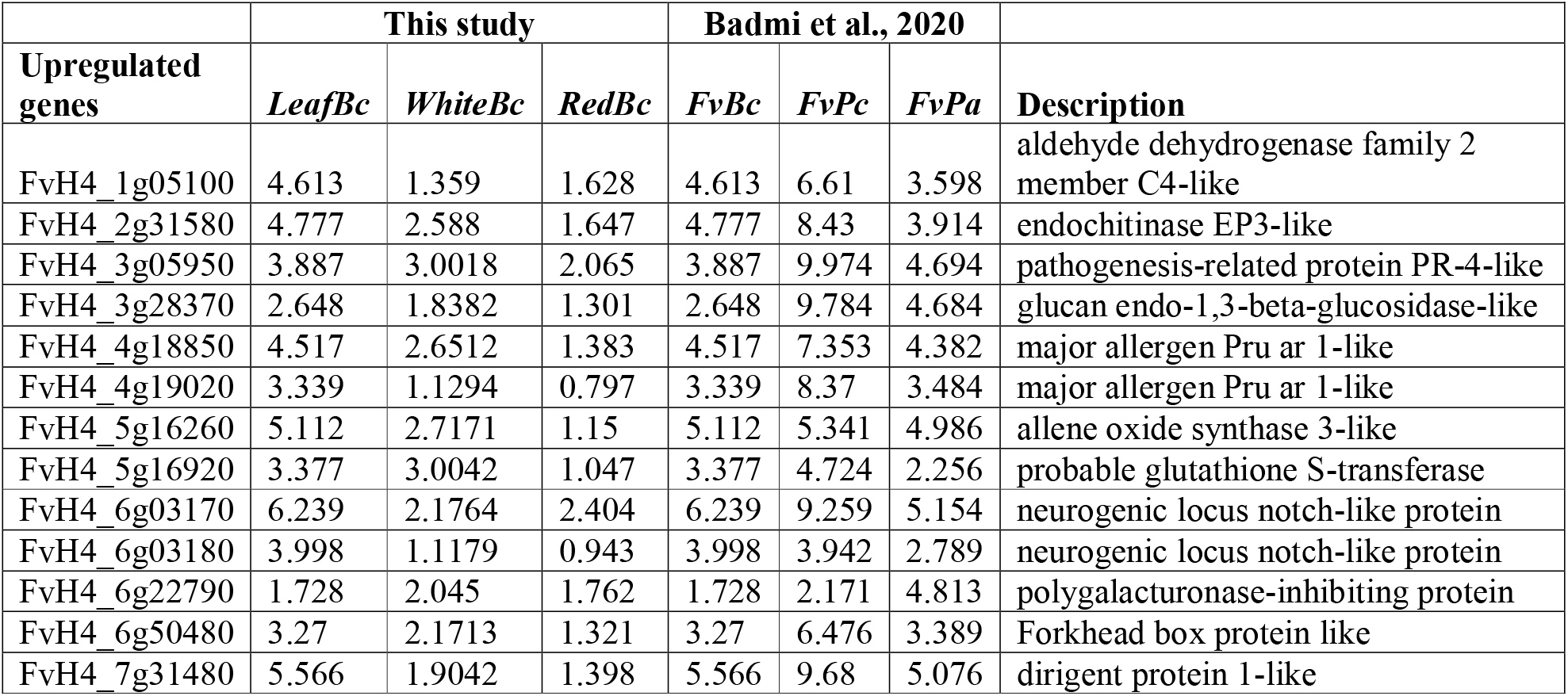
List of genes that are upregulated in three *F. vesca* tissues infected by *B. cinerea* (this study) as well as upon infection by three different pathogens in *F. vesca* (Badmi and Sheikh, 2020).

| Upregulated genes | This study |  |  | Badmi et al., 2020 |  |  | Description |
| --- | --- | --- | --- | --- | --- | --- | --- |
|  | *LeafBc* | *WhiteBc* | *RedBc* | *FvBc* | *FvPc* | *FvPa* |  |
| FvH4_1g05100 | 4.613 | 1.359 | 1.628 | 4.613 | 6.61 | 3.598 | aldehyde dehydrogenase family 2 member C4-like |
| FvH4_2g31580 | 4.777 | 2.588 | 1.647 | 4.777 | 8.43 | 3.914 | endochitinase EP3-like |
| FvH4_3g05950 | 3.887 | 3.0018 | 2.065 | 3.887 | 9.974 | 4.694 | pathogenesis-related protein PR-4-like |
| FvH4_3g28370 | 2.648 | 1.8382 | 1.301 | 2.648 | 9.784 | 4.684 | glucan endo-1,3-beta-glucosidase-like |
| FvH4_4g18850 | 4.517 | 2.6512 | 1.383 | 4.517 | 7.353 | 4.382 | major allergen Pru ar 1-like |
| FvH4_4g19020 | 3.339 | 1.1294 | 0.797 | 3.339 | 8.37 | 3.484 | major allergen Pru ar 1-like |
| FvH4_5g16260 | 5.112 | 2.7171 | 1.15 | 5.112 | 5.341 | 4.986 | allene oxide synthase 3-like |
| FvH4_5g16920 | 3.377 | 3.0042 | 1.047 | 3.377 | 4.724 | 2.256 | probable glutathione S-transferase |
| FvH4_6g03170 | 6.239 | 2.1764 | 2.404 | 6.239 | 9.259 | 5.154 | neurogenic locus notch-like protein |
| FvH4_6g03180 | 3.998 | 1.1179 | 0.943 | 3.998 | 3.942 | 2.789 | neurogenic locus notch-like protein |
| FvH4_6g22790 | 1.728 | 2.045 | 1.762 | 1.728 | 2.171 | 4.813 | polygalacturonase-inhibiting protein |
| FvH4_6g50480 | 3.27 | 2.1713 | 1.321 | 3.27 | 6.476 | 3.389 | Forkhead box protein like |
| FvH4_7g31480 | 5.566 | 1.9042 | 1.398 | 5.566 | 9.68 | 5.076 | dirigent protein 1-like |

## Author Contributions

RB conceived the idea, retrieved and analysed the data, wrote the manuscript.

